# Glycerol-driven Denitratation: Process Kinetics, Microbial Ecology, and Operational Controls

**DOI:** 10.1101/2021.09.25.461789

**Authors:** Matthew P. Baideme, Chenghua Long, Luke T. Plante, Jeffrey A. Starke, Michael A. Butkus, Kartik Chandran

## Abstract

Denitratation, the selective reduction of nitrate to nitrite, is a novel process when coupled with anaerobic ammonium oxidation (anammox) could achieve resource-efficient biological nitrogen removal of ammonium- and nitrate-laden waste streams. Using a fundamentally-based, first principles approach, this study optimized a stoichiometrically-limited, glycerol-driven denitratation process and characterized mechanisms supporting nitrite accumulation with results that aligned with expectations. Glycerol supported selective nitrate reduction to nitrite and near-complete nitrate conversion, indicating its viability in a denitratation system. Glycerol-supported specific rates of nitrate reduction (135.3 mg-N/g-VSS/h) were at least one order of magnitude greater than specific rates of nitrite reduction (14.9 mg-N/g-VSS/h), potentially resulting in transient nitrite accumulation and indicating glycerol’s superiority over other organic carbon sources in denitratation systems. pH and ORP inflection points in nitrogen transformation assays corresponded to maximum nitrite accumulation, indicating operational setpoints to prevent further nitrite reduction. Denitratation conditions supported enrichment of *Thauera* sp. as the dominant genus. Stoichiometric limitation of influent organic carbon, coupled with differential nitrate and nitrite reduction kinetics, optimized operational controls, and a distinctively enriched microbial ecology, was identified as causal in glycerol-driven denitratation.

## 1. Introduction

Conventional biological nitrogen removal (BNR), including energy and chemical-intensive nitrification and denitrification, is traditionally used to treat ammonium-laden (NH_4_^+^) waste streams. The advent of engineered processes that achieve oxidation of NH_4_^+^ to nitrite (NO_2_^-^), termed nitritation, combined with denitritation (reduction of NO_2_^-^ to nitrogen gas (N_2_)) or anaerobic ammonium oxidation (anammox) represent short-cut BNR alternatives to conventional BNR approaches. Such short-cut BNR processes can provide reductions in chemical (external carbon for denitrification and alkalinity for nitrification) and energy use (aeration for nitrification), driving the desire for NO_2_^-^ accumulation within these processes.

Alternatively, waste streams containing concomitantly high concentrations of NH_4_^+^ and nitrate (NO_3_^-^), such as those resulting from fertilizer^1^ and explosives manufacturing,^2,3^ provide similar energy and chemical reduction opportunities through distinct short-cut BNR processes. A particularly effective pathway for treating waste streams containing both NH_4_^+^ and NO_3_^-^ is through heterotrophic^4–9^ or autotrophic^10^ denitratation (selective reduction of NO_3_^-^ to NO_2_^-^) coupled with downstream anammox. A combined denitratation-anammox system used to treat waste streams containing equal concentrations of NH_4_^+^ and NO_3_^-^ would theoretically reduce aeration energy requirements by 100% and COD requirements by 80% compared to treatment of the same waste stream using conventional BNR. Recent studies^4–9^ on heterotrophic denitratation have focused on performance in lab-scale sequencing batch reactors (SBRs) driven by acetate, methanol, glucose, and sludge fermentation liquid due to the lack of sufficient readily biodegradable chemical oxygen demand (COD) in typical waste streams. These studies have primarily been observational in nature, with particular emphasis placed on empirically identifying parameters and conditions that potentially contributed to NO_2_^-^ accumulation, such as influent COD:N ratios, pH, ORP, and loading rates. Stoichiometric limitation of influent COD:N ratios, specifically, has been shown to influence endpoint nitrogen speciation.^11^ Various parameter combinations were optimized, denoted by the observation of stable NO_3_^-^-to-NO_2_^-^ conversion ratios as high as 90% during steady-state studies.^6^

The selection of an external COD source to drive denitrification is critical when attempting to maximize NO_2_^-^ accumulation. Traditionally, methanol has been one of the most widely used external COD sources for denitrification due to its low cost and wide availability.^12^ NO_2_^-^ accumulation has proven difficult with methanol due to methanol dehydrogenase’s direct delivery of electrons to cytochrome *c* and proximal to NO_2_^-^ reductase as opposed to distribution solely through the ubiquinol pool to NO_3_^-^ reductase similar to other carbon sources.^13–15^ The unique electron delivery locations during methanol oxidation within the respiratory denitrification chain potentially contribute to concomitant NO_3_^-^ and NO_2_^-^ reduction.

Several water resource recovery facilities are switching to glycerol due to the operational and safety risks associated with methanol.^12^ Glycerol is similar in cost to methanol and less expensive than ethanol and acetate,^16–18^ is available as a waste or byproduct,^19,20^ and has no known inhibitory effects on the anammox process, unlike methanol.^21^ NO_2_^-^ accumulation during glycerol supplementation was also anecdotally observed in full-scale treatment plants resulting in unintentional enrichment of anammox on the produced NO_2_^-^.^22^ Nevertheless, to fully realize the operating benefits that a denitratation-anammox system could offer, it is imperative for the parameters and conditions leading to NO_2_^-^ accumulation in a glycerol-driven denitratation system to be systematically identified, defined, and addressed in relation to reactor operating strategies.

Accordingly, the overarching goals of this study were to use a fundamentally-based, first principles approach to characterize the process kinetics, nitrogen conversion efficiencies, and microbial ecology of a glycerol-fed denitratation process, and identify concomitant reactor operating strategies. The specific objectives were to (1) control selective conversion of NO_3_^-^ to NO_2_^-^ through stoichiometric limitation of influent glycerol dose, (2) quantify the rates of NO_3_^-^ reduction relative to rates of NO_2_^-^ reduction and understand their impact on the selective accumulation of NO_2_^-^, (3) elucidate the microbial community structure under varied carbon-loading levels in a functional glycerol-driven denitratation process, and (4) identify operational controls and reactor operating strategies to maximize denitratation rates and efficiencies.

## 2. Materials and Methods

### 2.1. Experimental Set-up and Reactor Operation

A lab-scale SBR with a working volume, V=12 L, was operated at room temperature (22±2°C) for a period of 232 d. The SBR was operated at a hydraulic retention time (HRT) of 1 d, utilizing 4 cycles per day with each cycle consisting of a 90-min anoxic feed and react period, a 180-min anoxic react period, a 50-min settling period, and a 40-min decant period. SBR feed contained 100.0 mg/L NO_3_^-^-N as the terminal electron acceptor to simulate the influent of a high NO_3_^-^-containing waste stream typical of a fertilizer^1^ or explosives^2^ manufacturing facility, 25.0 mg/L NH_4_^+^-N (to support assimilation), and macro and trace nutrients (Table S1). pH was controlled automatically at 7.50±0.05 using 0.5 M HCl and 1.0 M NaHCO_3_ via a chemical dosing pump (Etatron D.S., Italy). Sludge wasting was controlled daily at the end of the anoxic feed and react period following COD exhaustion to maintain a solids retention time (SRT) of 3 d. Glycerol, diluted to a 15% solution by volume, served as the external COD source and was provided to meet influent COD:NO_3_^-^-N ratios from 2.4:1 to 5.0:1. Glycerol was fed at the end of the anoxic feed and react period so that examined influent COD:NO_3_^-^-N ratios were met during each cycle. Upon transitioning to each influent COD:NO_3_^-^-N ratio tested, a stabilization period of 4 x SRT was allowed prior to assessing performance relative to other conditions. Sequencing and timing of SBR cycles and daily solids wasting was controlled and maintained by peristaltic pumps (Masterflex, IL) using electronic timers (ChronTrol Corporation, CA).

### 2.2. Sample Collection and Wastewater Quality Analysis

All analytical procedures employed were in accordance with Standard Methods.^23^ Aqueous-phase samples were withdrawn during the decant period of the reactor cycle and concurrently from the influent for chemical species analysis after centrifugation (8,000 x G, 10 min, 4-8ºC) to remove cells and cell debris. NO_3_^-^ and NH_4_^+^ were measured using ion selective electrodes (Thermo Fisher Scientific, MA). NO_2_^-^ concentration was measured via diazotization and colorimetry.^23^ The fraction of influent NO_3_^-^ lost to nitrogenous gases was determined via mass balance on nitrogen. Centrifuged aqueous-phase samples were filtered using 0.20 μm syringe filters (A Chemtek, MA) and stored at -20°C. Dionex ICS-2100 ion chromatography using a Dionex IonPac AS-18 IC column (Thermo Fisher Scientific, MA) was used to confirm ion selective electrode and colorimetric measurements of NO_3_^-^ and NO_2_^-^ concentrations, respectively. Similarly, a Dionex IonPac AS-14 IC column (Thermo Fisher Scientific, MA) was used to quantify volatile fatty acid production during unbuffered *ex situ* batch kinetic assays. Separate aqueous-phase samples were extracted at the end of the anoxic react period and during the decant period of the reactor cycle to assess total biomass concentrations in the reactor and effluent, respectively, for SRT control. Aqueous-phase samples taken during the decant period were centrifuged (8,000 x G, 10 min, 4-8ºC) and filtered using 0.45 μm syringe filters (A Chemtek, MA) to assess remaining soluble COD (sCOD) concentrations (Hach Chemical Company, CO). Biomass concentrations were approximated by subtracting sCOD measurements from total COD measurements to determine particulate COD (pCOD) (Hach Chemical Company, CO).^24^ Additional aqueous-phase samples taken just prior to the end of the anoxic react period were centrifuged (8,000 x G, 10 min, 4-8ºC), supernatant was discarded, and cell pellets were preserved at -80°C for subsequent DNA extraction and 16S rRNA gene sequencing.

### 2.3. Feeding Strategy Experiments

Two feeding strategies were tested to maximize NO_2_^-^ accumulation. A semi-continuous feeding strategy delivered NO_3_^-^-containing SBR feed and glycerol continuously for the first 75 and 72 min, respectively, of the anoxic feed and react period (Fig. S1). A pulse feeding strategy delivered a pulse of NO_3_^-^-containing SBR feed and glycerol every 45 min for the first 270 min of the SBR cycle (Fig. S1). Feeding rates were controlled to maintain equivalent mass loading rates of NO_3_^-^ and glycerol and influent COD:NO_3_^-^-N ratios for the two feeding strategies.

### 2.4. Batch kinetic assays

Batch assays, *in situ* (within the SBR) and *ex situ*, were conducted to measure extant process kinetics and optimize operational controls, including batch duration, pH, and ORP. *In situ* assays followed previously described sampling collection and chemical analysis procedures. Aqueous-phase samples were obtained from the primary SBR at steady-state over the course of a single 360-min reactor cycle. *Ex situ* assays were carried out in an anoxic, sealed, spinner flask batch vessel with a working volume, V=1 L, at room temperature (22±2°C). Mixed liquor was taken from the primary SBR at steady-state during the feed and react period, washed 4 times using SBR feed without NO_3_^-^, and supernatant was discarded. Prior to extant kinetic batch assays, the medium was buffered to pH 7.50 using 0.5 M HCl and 1.0 M NaHCO_3_ and N_2_ gas was sparged until dissolved oxygen (DO) levels were equal to 0.01 mg/L O_2_, or the minimum practical limit of the InPro 6850i polarographic DO sensor with M300 transmitter (Mettler-Toledo, OH). pH was maintained at pH 7.50±0.05 by manual control. pH optimization batch assays were conducted within normal pH operating ranges (see Supporting Information (SI)). NO_3_^-^ and glycerol were dosed to meet the desired initial COD:NO_3_^-^-N ratio. NO_3_^-^ was dosed at the outset of the experiment (time=0 min) and the biomass was incubated for 30 min prior to the addition of glycerol to ensure that residual nitrogen species and glycerol from the primary SBR remaining in the washed mixed liquor were consumed prior to data collection. pH, ORP, and DO were measured and recorded continuously via an InPro 3253i/SG pH/ORP electrode and an InPro 6850i polarographic DO sensor, respectively, attached to an M300 transmitter (Mettler-Toledo, OH). Following extant kinetic batch assays, linear regression with R^2^≥95% of NO_x_-N species from time points of maximum concentration to minimum concentration for each respective species was performed with pCOD concentrations taken just prior to glycerol input to determine true specific rates of NO_3_^-^ reduction (sDNaR) (Eqn. 1) and NO_2_^-^ reduction (sDNiR) (Eqn. 2). NO_2_^-^ production resulting from NO_3_^-^ reduction was not accounted for in the determination of specific rates of NO_2_^-^ reduction, yet this remains representative of a true reduction rate. During the time points assessed for each influent COD:NO_3_^-^-N ratio, NO_3_^-^ removal was complete or near-complete (<3% of initial dose) except at influent COD:NO_3_^-^- N=2.5:1 where NO_3_^-^ concentration measurements confirmed no continued NO_3_^-^ reduction. pCOD measurements were used to determine maximum specific substrate consumption rates (Eqns. 1-2).

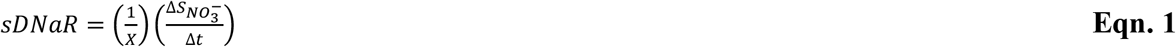

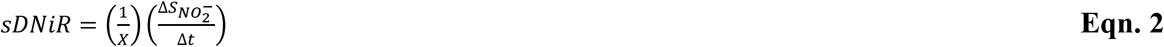

Where:

> *sDNaR*: maximum specific NO_3_^-^ consumption rate (mg NO_3_^-^-N/g VSS/h)
>
> *sDNiR*: maximum specific NO_2_^-^ consumption rate (mg NO_2_^-^-N/g VSS/h)
>
> *X*: volumetric biomass concentration approximated using pCOD measurements (g VSS/L)
>
> 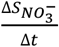 volumetric substrate (NO_3_^-^) consumption rate (mg NO_3_^-^-N/L/h)
>
> 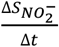 volumetric substrate (NO_2_^-^) consumption rate (mg NO_3_^-^-N/L/h)

### 2.5. DNA Extraction, Next-Generation Sequencing of Amplicon Library, and Bioinformatics

DNA was extracted from biomass samples and purified using a QIAamp DNA Mini Kit (Qiagen, Inc., MD). The quality and quantity of DNA were checked using a NanoDrop Lite spectrophotometer (Thermo Fisher Scientific, MA). Barcoded fusion primers with Ion Xpress™ sequencing adapters (Thermo Fisher Scientific, MA) and a 16S rRNA bacterial 1055F/1392R universal primer set were applied in each sample for multiplex sequencing. Amplification of genomic DNA targets was performed with iQ™ SYBR^®^ Green Supermix (Bio-Rad, CA) and purification via Agencourt AMPure XP Reagent (Beckman Coulter, CA). Library quantification was performed with an Agilent DNA 1000 Kit (Agilent, CA). Template preparation with the DNA library followed by Ion Spheres Particle (ISP) enrichment was performed using Ion OneTouch2 (Ion PGM Hi-Q View OT2 Kit). Enriched ISP was loaded onto an Ion Torrent 318 v2 BC chip and run on an Ion Torrent Personal Genome Machine (Ion PGM Hi-Q View Sequencing Kit). Ion Torrent Suite software was used for base calling, signal processing, and quality filtering (Phred score of >15) of the raw sequences. The 1055F/1392R universal primer set targeted sequences of approximately 350 base pairs (bp). Mothur software was used to initially screen out likely incorrect amplicon sequences with bp lengths more than 50 bp different than the target sequence length.^25^ AfterQC software was utilized to further delete bad quality reads (Phred score of <20) and trim the tails of reads where quality dropped significantly.^26^ DADA2 programming via R Studio software was used to produce a table of non-chimeric amplicon sequence variants from the demultiplexed fastq files.^27^ QIIME2 software was applied in conjunction with the Silva version 132 reference taxonomy for further post-sequencing bioinformatic analysis.^28^

### 2.6. Nitrogen Conversion Calculations

Reactor performance was normalized with respect to the influent characteristics. A NO_2_^-^ accumulation ratio (NAR) (Eqn. 3) was defined to relate the accumulation of NO_2_^-^ to the removal of NO_3_^-^.^29^ A NAR equal to 100% indicated that all NO_3_^-^ removed accumulated as NO_2_^-^ compared to terminal reduction to N_2_ gas, for which the NAR would be 0%.

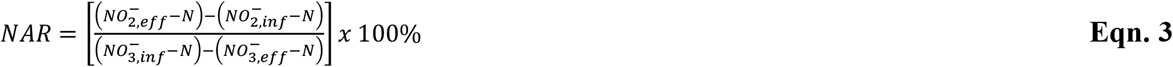

NO_3_^-^ reduction was also classified in terms of a NO_3_^-^ reduction ratio (NRR) (Eqn. 4), which normalized the conversion of NO_3_^-^ to the influent NO_3_^-^ concentration.^9^ A NRR equal to 100% would indicate conversion of all influent NO_3_^-^ to any reduced form, while a NRR of 0% would indicate no conversion.

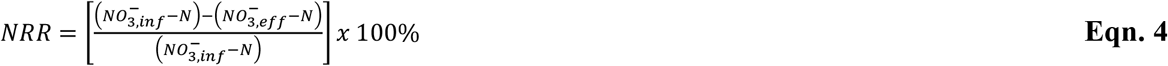

## 3. Results and Discussion

### 3.1. Denitratation Reactor Performance

The influent COD:NO_3_^-^-N ratio required for glycerol-driven denitrification (NO_3_^-^-N to N_2_ reduction) was thermodynamically^30^ determined to be 5.9:1 (see SI). This corresponded well with experimentally-determined operational ratios of 4.2:1 to 5.6:1,^16,20,31^ although the lowest reported ratio^16^ may not be fully representative as it was determined via *ex situ* batch assays as opposed to steady-state continuous flow bioreactor or SBR operation. Stoichiometric analysis revealed that influent COD:NO_3_^-^-N=2.4:1 (see SI) would provide only enough electrons via COD oxidation to reduce NO_3_^-^ to NO_2_^-^ on a theoretical electron equivalence basis as opposed to full denitrification. Therefore, influent COD:NO_3_^-^-N ratios between 2.4:1 and 5.9:1 were referred to as stoichiometrically-limited for the purposes of this study. These calculations form the fundamentally-based foundation to the first principles approach used in this study to conduct and interpret the results of glycerol-driven denitratation presented herein.

The utilization of glycerol as the external COD source and electron donor resulted in significant NO_2_^-^ accumulation at stoichiometrically-limited influent COD:NO_3_^-^-N ratios from 2.5:1 to 5.0:1, indicating that the use of glycerol was feasible to sustain a denitratation process. The highest degrees of NO_3_^-^ removal and NO_2_^-^ accumulation, as a function of influent COD:NO_3_^-^-N ratio during steady-state SBR operation, occurred at influent COD:NO_3_^-^-N=3.0:1 (Fig. 1). This resulted in an average NO_2_^-^ accumulation of 60.8±11.5 mg/L NO_2_^-^-N (n=10) and NAR of 62%, indicating that 62% of the NO_3_^-^ reduced was converted to NO_2_^-^ rather than terminally reduced to N_2_ gas. Additionally, the NRR was determined to be 96%, indicating that a majority of the influent NO_3_^-^ was converted leaving only approximately 4% of influent NO_3_^-^ in the effluent (Table 1). Accumulation of NO_2_^-^ at influent COD:NO_3_^-^-N=2.8:1 compared to influent COD:NO_3_^-^-N=3.0:1 was not significantly different (p=0.49, α=0.05, n=10). Substantial NO_3_^-^ accumulation occurred at influent COD:NO_3_^-^-N=2.8:1 (31.7±11.4 mg/L NO_3_^-^-N, n=11), signifying that this ratio was less operationally optimal compared to influent COD:NO_3_^-^- N=3.0:1. The observed NO_3_^-^ accumulation at influent COD:NO_3_^-^-N=2.5:1 and 2.8:1 may be due to lower COD-supported biomass concentrations leading to reduced denitrification rates. However, effluent sCOD concentrations were negligible signifying that glycerol was nearly completely consumed (sCOD and biomass concentration data not shown). *In situ* performance profiles (Fig. 2) did not show significant endogenous denitrification, potentially indicating that COD uptake and storage was minimal. Rather, the observed NO_3_^-^ accumulation in these cases indicated that the influent COD:NO_3_^-^-N was not sufficient,^9^ potentially due to unrealized COD requirements for cell maintenance and synthesis^32^ or additional demand by fully denitrifying microorganisms remaining in the microbial community. Therefore, influent COD:NO_3_^-^-N=3.0:1 was selected as the optimal ratio due to the similar NO_2_^-^ accumulation to influent COD:NO_3_^-^- N=2.8:1 coupled to less than 4% of the influent NO_3_^-^ remaining in the effluent. The high sensitivity at influent COD:NO_3_^-^-N<3.0:1 highlighted significant implication for accurate system operation and control. A minimal reduction in influent COD:NO_3_^-^-N ratio from 3.0:1 to 2.8:1 yielded a sevenfold increase in effluent NO_3_^-^, signifying that strict control of the glycerol-driven denitratation system must be maintained. To this end, online dosing control^17^ based on appropriate signals of reactor performance seems necessary to maximize concomitant NO_3_^-^-N conversion selectively to NO_2_^-^ during partial denitratation.

**Table 1.**
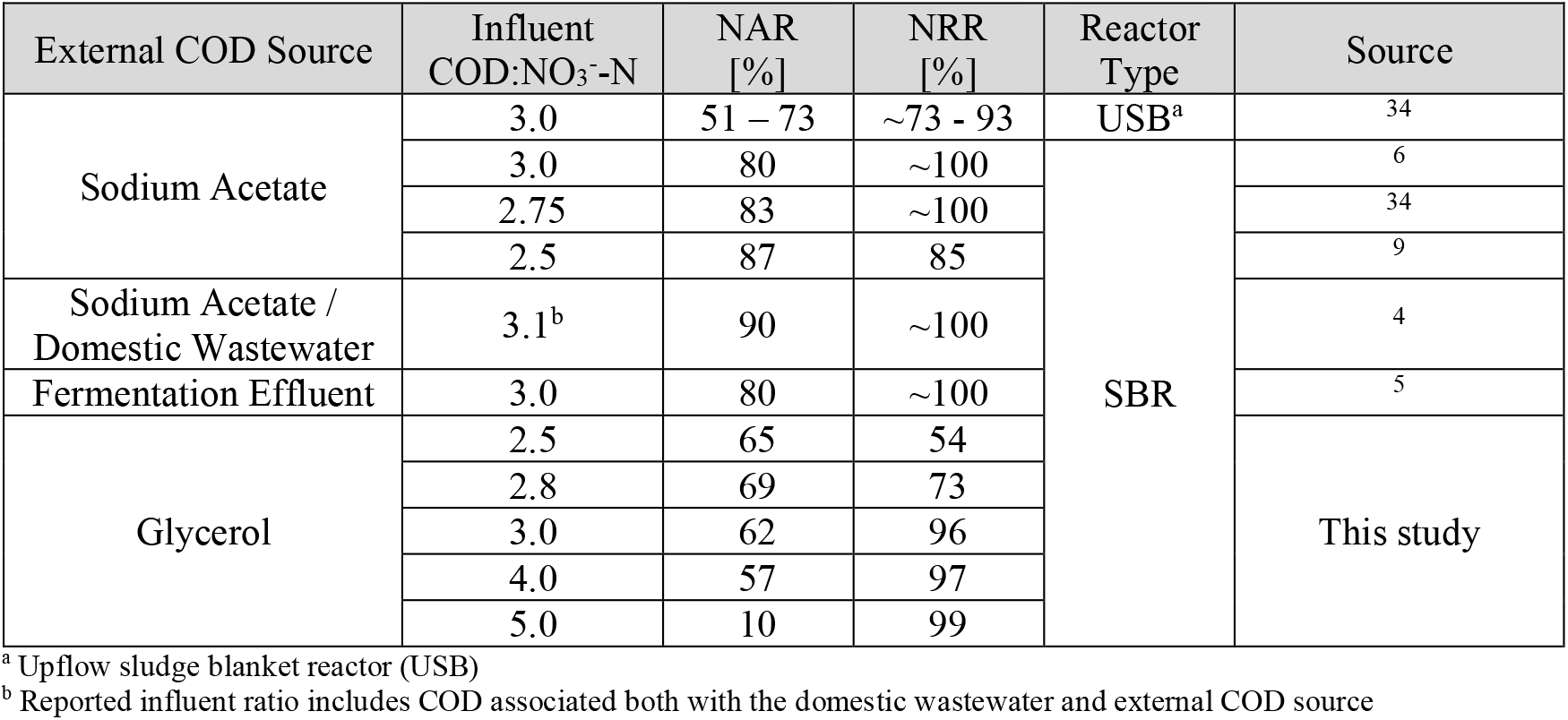
Influence of external COD source and influent COD:NO_3_^-^-N ratios on denitratation performance.

**Figure 1.**
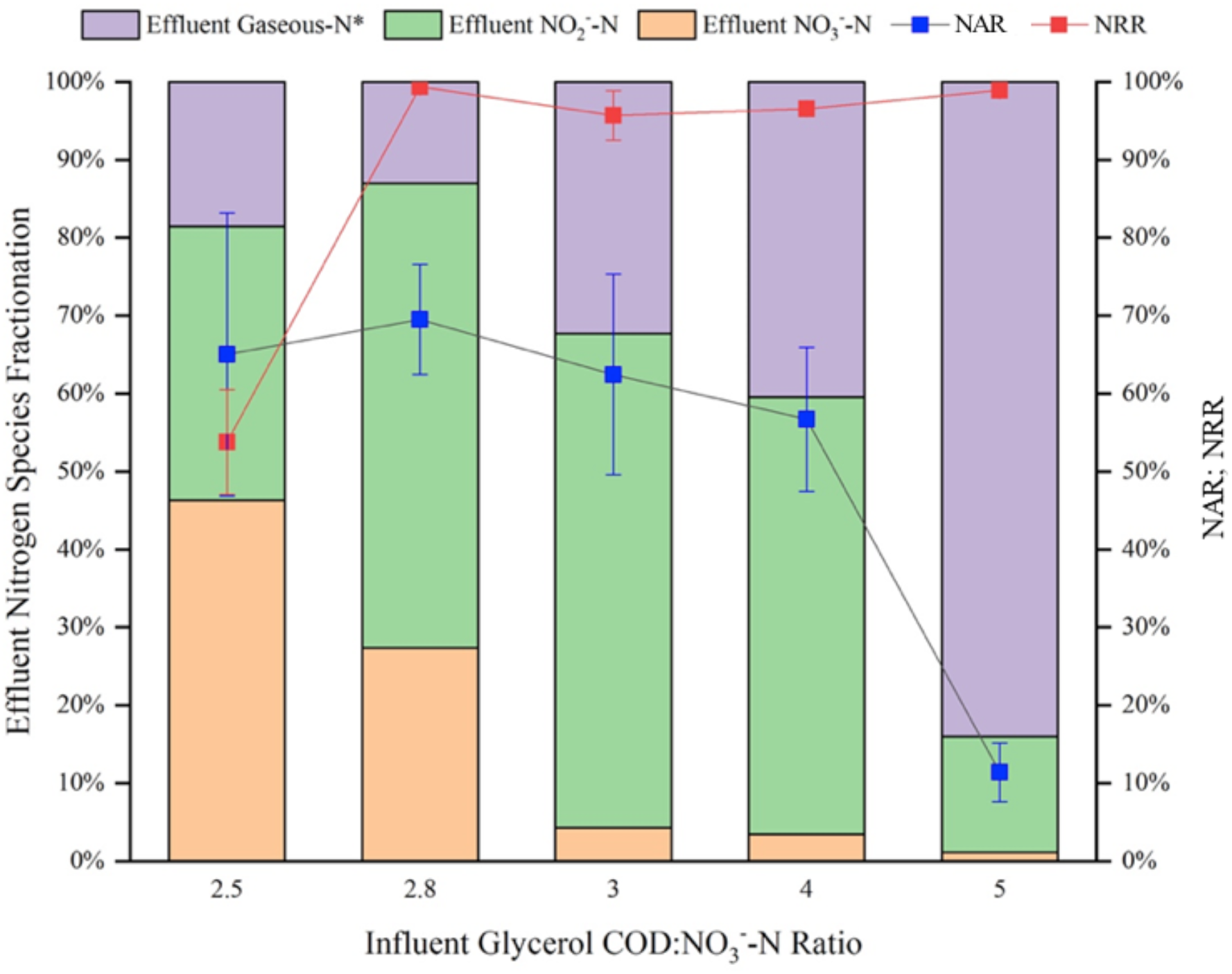
Steady-state denitratation performance and respective NAR and NRR assessed at each influent COD:NO_3_^-^-N ratio. ^*^Effluent gaseous-N contributions were calculated via mass balance

**Figure 2.**
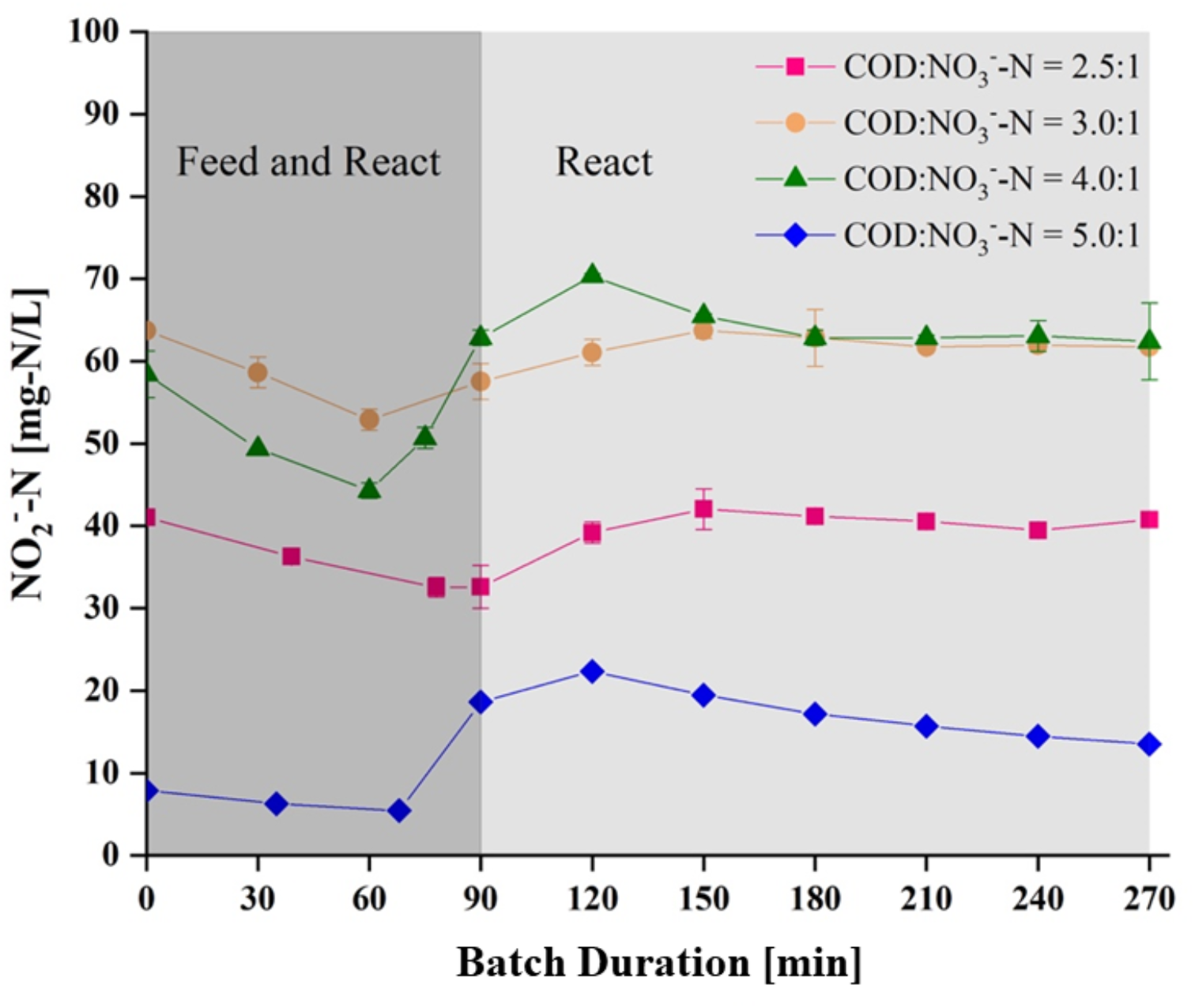
Representative in situ NO_2_^-^-N profiles identified the optimal batch duration obtained during steady-state operation at each respective influent COD:NO_3_^-^-N ratio. Optimal batch durations corresponded to the points of maximum NO_2_^-^ accumulation at each respective influent COD:NO_3_^-^-N ratio. Decreases in NO_2_^-^ concentrations during the feed and react period were attributed to dilution.

Analysis of variance (ANOVA) across the influent COD:NO_3_^-^-N ratios identified a statistically significant difference in NAR (p=4.8×10^−11^, α=0.05, n=38) with a decrease from 62% to 11% as the influent COD:NO_3_^-^-N ratio approached that for glycerol-driven denitrification (5.9:1; see SI). Further Holm-Sidak post-hoc multiple comparison analysis indicated that the significant difference in NAR was primarily caused by the expectedly lower NAR at influent COD:NO_3_^-^-N=5.0:1 (p<9.7×10^−5^ for all comparisons, α=0.05; Table S2). The decrease in NAR from influent COD:NO_3_^-^-N=4.0:1 to 5.0:1 was most likely attributable to excess available COD.

Previous studies^4,6^ observed that varying the influent COD:NO_3_^-^-N ratio had a negligible effect on the NAR determined at the point of maximum NO_2_^-^ accumulation during *ex situ* batch experiments, while a separate batch study^33^ concluded that the COD source, as opposed to the influent COD:NO_3_^-^-N ratio, impacted the NAR more readily. In contrast, another separate batch study^7^ concluded that NO_2_^-^ accumulation was influenced by both the COD source and COD dosing. While insightful, the utility of these results^4,6,7^ to guide steady-state denitratation processes is limited as these studies failed to acclimate their batch experiment seed sludge to the conditions being investigated, which likely contributed to their discrepancy with the current study. Despite investigating the impact of various influent COD:NO_3_^-^-N ratios, Ge et al.^7^ utilized a fully denitrifying inoculum, whereas Du et al.^6^ inoculated batch experiments assessing various influent COD:NO_3_^-^-N ratios with a microbial community acclimated to a single stoichiometrically-limited influent COD:NO_3_^-^-N ratio. Both seed sludges likely contained phenotypes with NO_2_^-^ accumulation capabilities different than those expected following acclimation to the investigated conditions. Cao et al.^4^ did not report conditions of their batch inoculum.

In an improvement over these previous efforts, our current study utilized a sludge stabilization and acclimation period of 4 x SRT following influent COD:NO_3_^-^-N ratio changes. This intentionally allowed the microbial community to adapt to the influent COD:NO_3_^-^-N ratio being investigated. In doing so, it was observed that the influent COD:NO_3_^-^-N ratio had similar impacts on NAR during both steady-state operation (Fig. 1) and *ex situ* batch assays, with NO_2_^-^ accumulation decreasing as influent COD:NO_3_^-^-N ratios increased (Fig. S2).

In comparison to other steady-state operation studies^6,9,34^ using primarily sodium acetate as the external COD source, glycerol-driven NARs were at least 10% lower (Table 1). While most reported acetate-driven denitratation NARs were greater than 80%, glycerol-driven denitratation yielded NARs less than 70%. These respective acetate-driven steady-state studies^6,9,34^ were deemed reasonable comparisons due to similar COD dosing regimens and results were reported for study periods sufficient in length to assume microbial community acclimation to and stabilization at the studied conditions. Despite this, the assessment of reactor performance based solely upon reported NARs can be misleading as the index does not account for complete or other conversion of influent NO_3_^-^. Thus, NAR=100% does not necessarily indicate that all influent NO_3_^-^ was converted. Several studies,^4–6,34^ however, reported NRRs of nearly 100% that when coupled with a NAR approaching 100% indicated near-perfect denitratation performance (Table 1). It follows then that optimal performance in the current study occurred at influent COD:NO_3_^-^-N=3.0:1 with NAR=62% and NRR=96%. The inability of glycerol to achieve similar efficiency to acetate- or fermentate-driven denitratation is not currently understood. Possible explanations include a greater intracellular carbon and microbial energy storage mechanism during low substrate availability,^35,36^ the COD-source supported enrichment of a microbial consortium with a greater abundance of true denitrifiers,^37^ an inefficient metabolism in support of denitratation due to a less direct assimilability of glycerol, or the downstream delivery of electrons on the electron transport chain similar to methanol.^14,15^

Effluent sCOD measurements, as an estimation of residual glycerol concentration, averaged 9.4±8.8 mg/L COD (n=29) across all influent COD:NO_3_^-^-N ratios assessed. The *ca*. 96% average decrease from influent to effluent sCOD indicates that nearly all of the glycerol was consumed, and that reactor cycle duration was adequate for COD consumption.

A likely contributing factor to the need for a higher than the theoretical influent COD:NO_3_^-^-N ratio (see SI) was an incomplete enrichment for a solely denitratating or progressive onset^38^ phenotype-dominated microbial community. The presence of microorganisms that express a complete denitrification metabolic pathway or those that exhibit a rapid, complete onset of denitrification genes^38^ would impose a competitive demand on influent COD, thus decreasing its availability for selective reduction of NO_3_^-^ to NO_2_^-^. This additional COD demand would result in a high NRR but low NAR, or significant gaseous-N products with limited NO_2_^-^ accumulation, which was supported by the results herein (Table 1).

### 3.2. Process Kinetics

Notably, extant kinetic analysis indicated that transient NO_2_^-^ accumulation at all influent COD:NO_3_^-^-N ratios assessed was potentially due to at least one order of magnitude greater specific rates of NO_3_^-^ reduction compared to the specific rates of NO_2_^-^ reduction driven by glycerol (Table 2).^39^ Observed performance at influent COD:NO_3_^-^-N>3.0:1 (Fig. S2) also supported this assertion as the maximum NO_2_^-^ accumulated never equaled the initial NO_3_^-^ concentration, indicating that there was concomitant reduction of NO_3_^-^ and NO_2_^-^. However, performance at influent COD:NO_3_^-^-N=3.0:1 resulted in near-complete selective reduction of NO_3_^-^ to NO_2_^-^ prior to terminal reduction to N_2_ gas (Fig. S2). It should be emphasized that the kinetic profiles in Fig. S2 were obtained from acclimated biomass from individual SBRs operated for at least 4 x SRT at each influent COD:NO_3_^-^-N ratio.

**Table 2.**
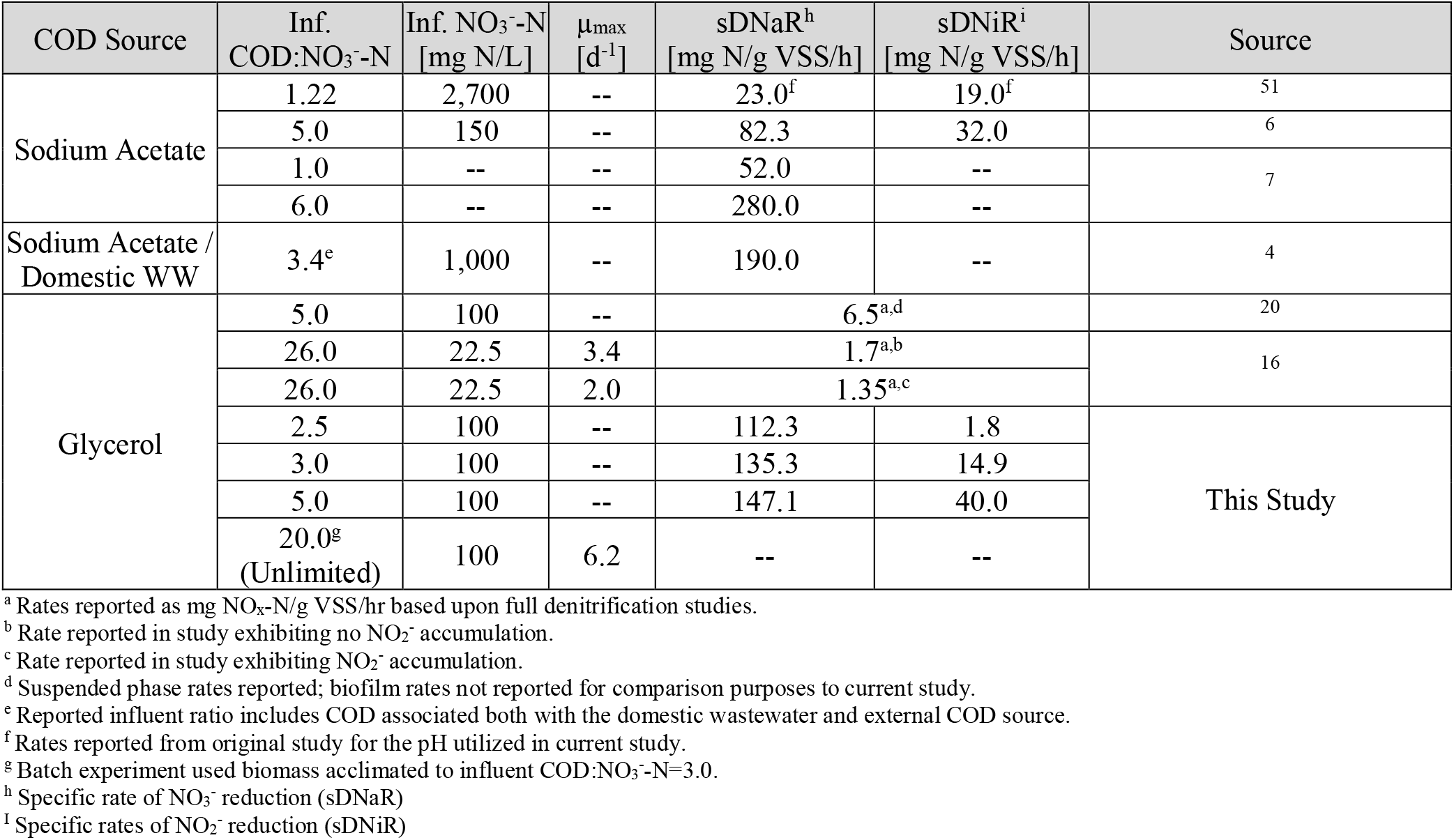
Summary of process kinetic parameters for both full denitrification and denitratation studies with respect to external COD source and influent COD:NO_3_^-^-N ratio.

In general, measured specific rates of NO_3_^-^ reduction and μ_max_ values were higher than those previously reported for glycerol-driven full denitrification studies (Table 2) and may be due to differences in the microbial community that was selected for by stoichiometric limitation during our current denitratation-specific study. Glycerol-driven specific rates of NO_3_^-^ reduction values were nearly double those reported for acetate-driven systems at similar influent COD:NO_3_^-^-N ratios, but slightly lower than those observed in an experiment utilizing a combination of external COD sources garnered from sodium acetate and endogenous carbon in a domestic wastewater stream (Table 2). The ratios of sDNaR:sDNiR achieved in this study with glycerol across different influent COD:NO_3_^-^-N values were also higher than previously reported with acetate (Table 2). This difference may be due to variations in the direct assimilability of each COD source with more assimilable COD sources such as glycerol or endogenous carbon in these cases supporting greater specific rates of NO_3_^-^ reduction,^32^ or the COD source-supported microbial community.

### 3.3. NO_2_^-^ Accumulation through the Management of Operational Controls

#### 3.3.1. Denitratation Control via Batch Duration

Batch duration was identified as an effective process control parameter to maximize NO_2_^-^ accumulation. The duration of the anoxic feed and react period could be shortened to achieve comparable or improved performance. NO_2_^-^ concentrations decreased following peaks of NO_2_^-^ accumulation at higher influent COD:NO_3_^-^-N ratios (4.0:1, 5.0:1; Fig. 2). This decrease was not observed at influent COD:NO_3_^-^-N=3.0:1, indicating that excess COD remained following completion of denitratation at higher ratios. Despite minimal NO_2_^-^ reduction following peak NO_2_^-^ accumulation at influent COD:NO_3_^-^-N=2.5:1, overall performance remained low, making this ratio less effective at achieving partial denitratation (Table 1; Fig. 2).

Results generally supported that influent COD:NO_3_^-^-N ratios have an inverse relationship with time to maximum NO_2_^-^ accumulation during the anoxic react period. Batch duration could be reduced to 150 minutes or less, or the time to maximum NO_2_^-^ accumulation (Fig. 2). Subtraction of the feed and react period of the SBR cycle from the reduced batch duration, by extension, would yield an optimal react time equivalent to a continuous flow system’s HRT (Fig. 2). The optimal react time is representative of when glycerol is available for NO_3_^-^ reduction in both systems. Therefore, the identified optimal react times in our SBR system would be equivalent to HRTs of approximately 30 minutes (COD:NO_3_^-^-N=4.0:1 and 5.0:1) to 60 minutes (COD:NO_3_^-^-N=2.5:1 and 3.0:1) in continuous flow systems operating at each respective influent COD:NO_3_^-^-N ratio.

#### 3.3.2. Denitratation Control via pH and ORP

During unbuffered and non-carbon limited operation (influent COD:NO_3_^-^-N≥5.9:1), the denitratation-dominated phase of the denitrification profile exhibited a distinct decrease in the reactor’s pH and increase in the ORP until both reached inflection points after which pH increased and ORP decreased (Fig. 3). At this inflection point, NO_3_^-^ reduction decelerated due to the depletion of available NO_3_^-^ allowing for observable concomitant NO_2_^-^ reduction thus decreasing the NAR and negatively impacting the objective of maximizing NO_2_^-^ accumulation. Continuous monitoring of pH and ORP could provide an observable real-time control to maximize denitratation. While feedforward online control of COD dosing tied to influent NO_x_ loading has proven effective in controlling denitratation,^17^ this system requires online NO_x_ sensors which may not be achievable at all plants due to potentially high capital^40^ and maintenance costs.^41^ Rather, denitratation control via pH and ORP observation could provide a backup check or serve as a less costly alternative^40^ with widely available and utilized sensors.

**Figure 3.**
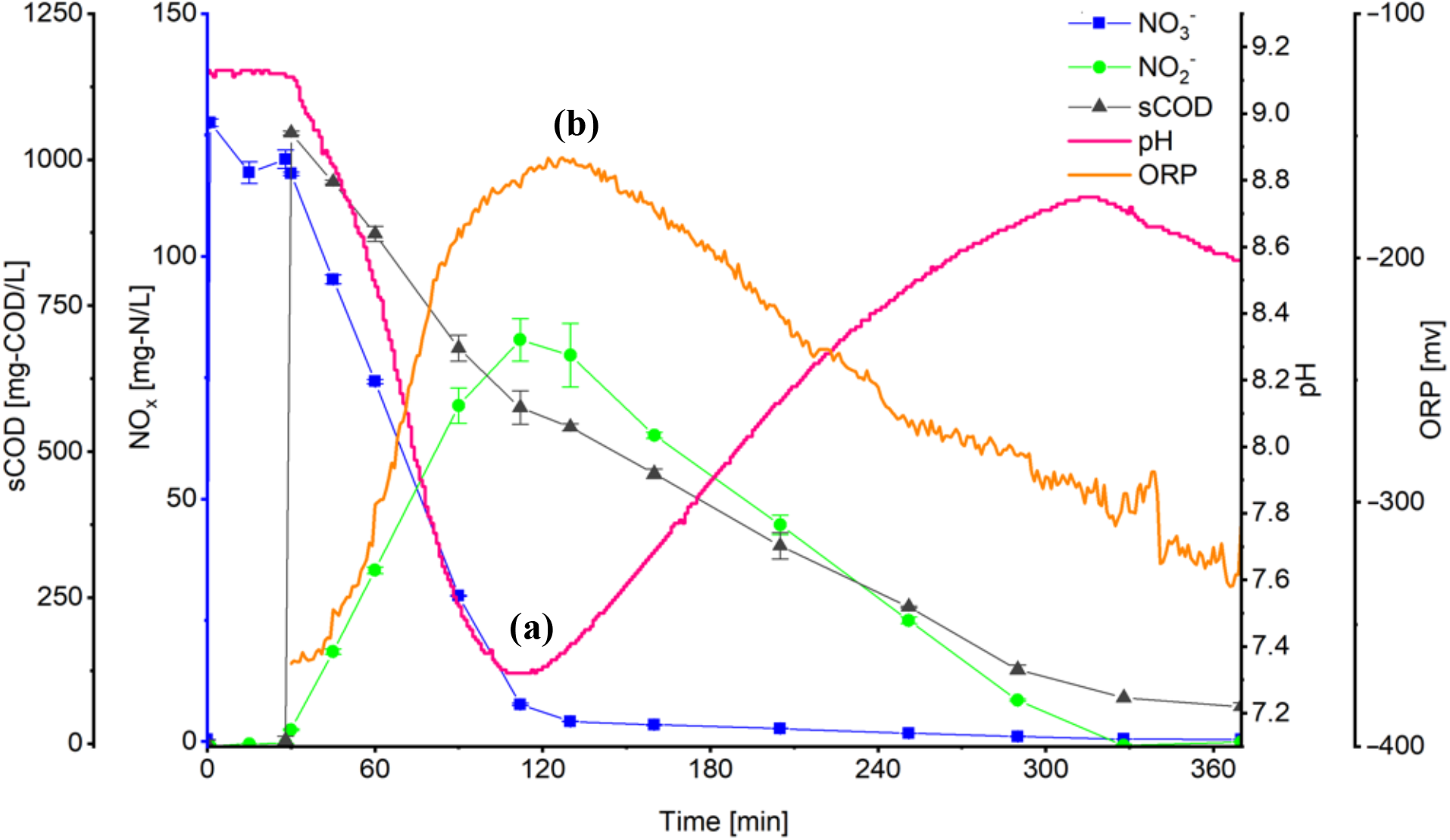
NO_x_, pH, and ORP profiles depicting the pH (a) and ORP (b) inflection points at the point of maximum NO_2_^-^ accumulation prior to which denitratation was dominant and after which denitritation became dominant (influent COD:NO_3_ ^-^-N=10.0:1; microbial ecology acclimated to influent COD:NO_3_^-^-N=3.0:1). Influent COD was provided in excess and beyond that at which biomass was acclimated in order to drive the process beyond denitratation and demonstrate the ability of pH and ORP to serve as denitratation process controls even under non-ideal influent COD:NO_3_^-^-N ratios.

pH and ORP were previously reported as control parameters for denitrification driven by acetate, methanol, endogenous carbon, soybean wastewater, and brewery wastewater.^6,7,33,42,43^ Contrary to the distinct glycerol-driven pH and ORP profile observed in the current study, Ge et al.^7^ and Du et al.^6^ described acetate-driven profiles exhibiting a general increase in pH whereby a “turning point” separated denitratation from denitritation. However, the observed pH profiles obtained experimentally in our study (Fig. 3) are in excellent concurrence with theoretically calculated net <u>production</u> of 0.43 equivalents of acidity per mole NO_3_^-^ reduced to NO_2_^-^ (Eqn. 5), which supported the observed pH fluctuation profiles.

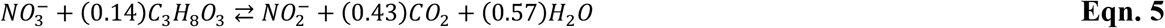

For completeness, stoichiometry (Eqn. 6) reveals that denitritation should result in a net <u>consumption</u> of 0.36 equivalents of acidity per mole NO_2_^-^ reduced to N_2_ gas at pH 7.5.

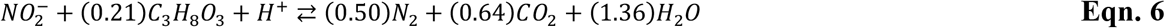

#### 3.3.3. Denitratation Control via Feeding Strategy

The pulse feeding strategy resulted in a statistically significant improvement in denitratation performance (α=0.05; n=8) over the semi-continuous feeding strategy in both NO_2_^-^ accumulation (p=0.03) and NO_3_^-^ reduction (p=0.0003), indicating that feeding methodology impacted the performance of the system (Table S3). As both feeding strategies maintained equivalent influent COD:NO_3_^-^-N ratio per substrate pulse or for the duration of the semi-continuous feeding period, this difference in system performance was thought to be influenced by the temporal distribution of substrate pulses. Those pulses occurring later in the anoxic feed and react period may have limited the time for the biotransformation of NO_3_^-^ to gaseous nitrogen thus allowing for greater NO_2_^-^ accumulation. This is counter to the semi-continuous feeding strategy, where fully denitrifying microorganisms within the microbial community had the full anoxic feed and react period to reduce influent NO_3_^-^. Therefore, in a continuous-flow BNR process, the spatial location of introducing glycerol could be another factor to promote partial denitratation if possible. Optimizing the dosing location of electron donors is quite widely practiced for increasing the efficiency of COD utilization even for full denitrification in step-feed BNR or Bardenpho configurations.^44^

### 3.4. Microbial Ecology

*Proteobacteria* was the most dominant phylum out of 14 identified at all influent COD:NO_3_^-^-N ratios (Fig. 4a). *β-Proteobacteria* made up at least 73% of the *Proteobacteria* phylum at all influent COD:NO_3_^-^-N ratios. In a survey of wastewater denitrifying bacterial 16S rDNA sequences retrieved from GenBank, Lu et al.^45^ found that approximately 72% of prokaryotic microorganisms displaying denitrifying capabilities were taxonomically affiliated with *Proteobacteria*, while *β* sub-class affiliated microorganisms were typically abundant in denitrifying activated sludge,^1,45,46^ similar to the findings herein.

**Figure 4.**
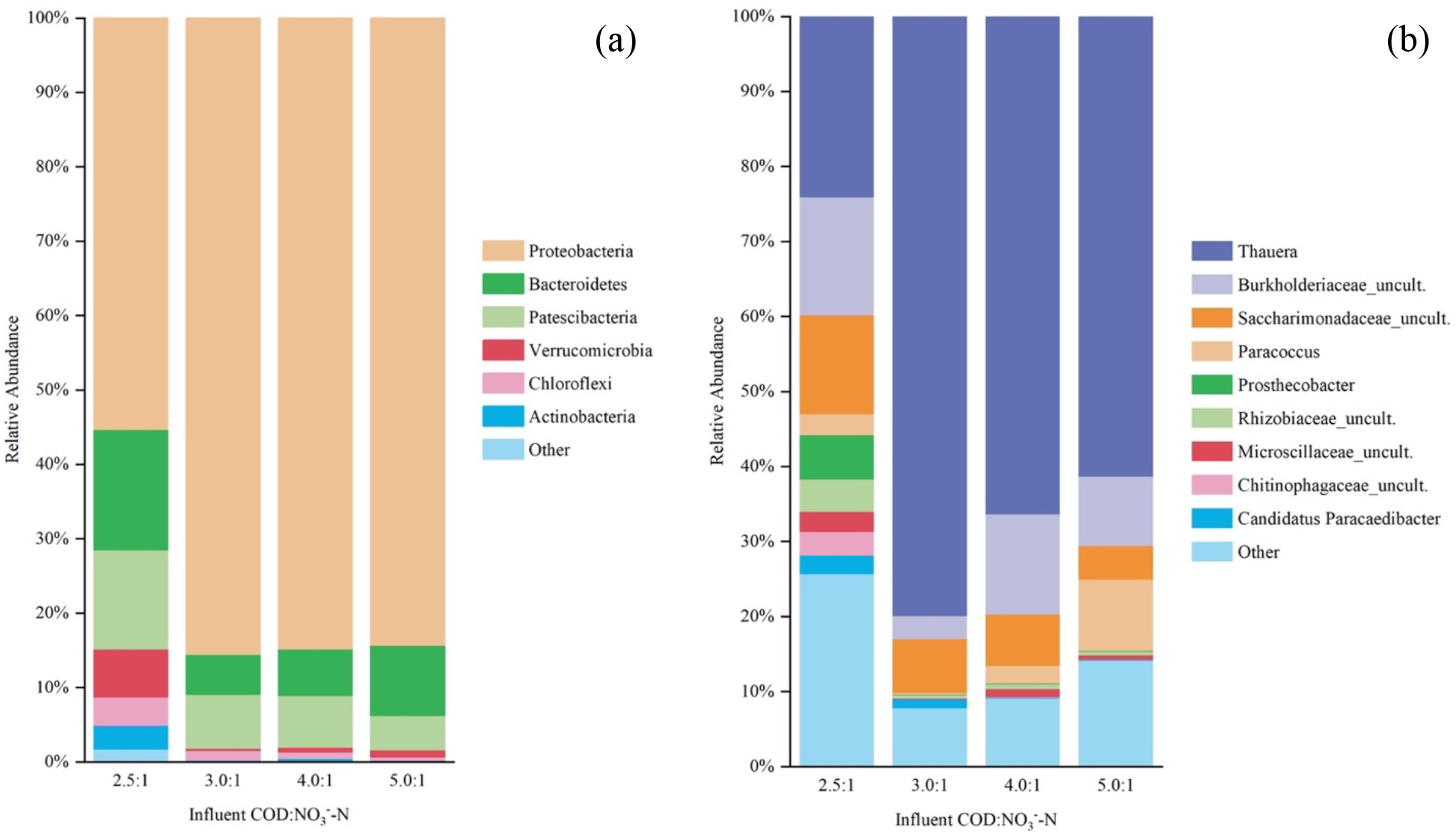
Taxonomic analysis of the microbial consortium at the phylum (a) and genus (b) taxonomic levels under optimal operating conditions (influent COD:NO_3_^-^-N=3.0:1, SRT=3 d). The grouping “Other” comprises OTUs with less than 1% total relative abundance (among all samples summed).

Within *β-Proteobacteria*, the *Rhodocyclaceae* and *Comamonadaceae* families were identified as those mainly involved in denitrification in activated sludge.^46,47^ Our findings supported this as *Thauera* sp., that belongs to the *Rhodocyclaceae* family within *β-Proteobacteria* was enriched as the most dominant genus with a relative abundance of nearly 80% at influent COD:NO_3_^-^-N=3.0:1 (Fig. 4b). *Comamonadaceae* fam. was not found, indicating that their enrichment may not be favored under stoichiometrically-limited conditions imposed herein. Certain *Thauera* spp. strains were characterized according to two distinct regulatory phenotypes,^48^ including the immediate and simultaneous onset of all denitrification genes with no detectable NO_2_^-^ accumulation, as well as the progressive and sequential onset of denitrification cascade genes with appreciable NO_2_^-^ accumulation.^38^ Selective pressures were not identified for either, although the selection for progressive onset denitrifiers would be critical to facilitate denitratation. The coupling of a high relative abundance of *Thauera* sp. (Fig. 4b), high NRR, and high NAR (Table 1), with the ability to perform full denitrification when presented with sufficient COD (Fig. S2) indicated that the application of stoichiometric limitation in the influent COD:NO_3_^-^-N as a selective pressure may favor the progressive onset over rapid, complete onset phenotype. *Thauera* sp. may represent a key functional microorganism for denitratation systems indicated by its decreasing relative abundances away from the optimal influent COD:NO_3_^-^-N (Fig. 4b). Several recent denitratation-specific studies^4,6,9,49^ further supported this argument with reported *Thauera* sp. relative abundances from 55% to 73% under limited influent COD:NO_3_^-^-N ratios with acetate as the external COD source despite different seed sludges. In comparison, acetate-driven full denitrification studies reported no more than 12% relative abundance of *Thauera* sp.^46,50^ Therefore, the application of a stoichiometrically-limited influent COD:NO_3_^-^-N ratio as a selective pressure in a denitratation system may impart a stronger impact on the denitrifying community structure than previously recognized.

## 4. Conclusions

Denitratation, with downstream anammox processes, offers chemical and energy reductions through resource-efficient BNR of NH_4_^+^ and NO_3_^-^-laden waste streams. A fundamentally-based, first-principles approach was used to propose an influent COD:N ratio and other operating parameters that would promote denitratation and experimental results aligned with expectations. Glycerol supported the process kinetics and microbial ecology necessary to selectively convert NO_3_^-^ to NO_2_^-^ in denitratation systems. Process control strategies, including influent COD loading and pH, ORP, and batch duration operational setpoints were identified and used to further define reactor operating strategies that could maximize denitratation performance. Significant enrichment indicated *Thauera* sp. may represent a key functional microorganism in denitratation systems. This study implicated stoichiometric limitation of influent organic carbon, unique microbial community enrichment, and differential NO_3_^-^ and NO_2_^-^ reduction kinetics as determinant factors in glycerol-driven denitratation.

## ADDITIONAL INFORMATION

E-supplementary data can be found in online version of the paper.

## AUTHOR INFORMATION

### Author Contributions

The manuscript was written through contributions of all authors. All authors gave approval to the final version of the manuscript. All authors contributed equally.

### Notes

The authors declare no competing financial interests.

## ACKNOWLEDGMENTS

This study was supported by Project Director, Joint Services, project USMA1740. Views and opinions expressed or implied herein are solely those of the authors and should not be construed as policy or carrying the official sanction of the Department of Defense, United States Army, United States Military Academy, or other agencies or departments of the U.S. Government.

## REFERENCES

1 C. S. Srinandan, M. Shah, B. Patel and A. S. Nerurkar, Bioresour. Technol., 2011, 102, 9481–9489.

2 P. Cyplik, R. Marecik, A. Piotrowska-Cyplik, A. Olejnik, A. Drożdżyńska and Ł. Chrzanowski, Water, Air, Soil Pollut., 2012, 223, 1791–1800.

3 J. Shen, R. He, W. Han, X. Sun, J. Li and L. Wang, J. Hazard. Mater., 2009, 172, 595–600.

4 S. Cao, R. Du, B. Li, S. Wang, N. Ren and Y. Peng, Chem. Eng. J., 2017, 326, 1186–1196.

5 S. Cao, S. Wang, Y. Peng, C. Wu, R. Du, L. Gong and B. Ma, Bioresour. Technol., 2013, 149, 570–574.

6 R. Du, Y. Peng, S. Cao, B. Li, S. Wang and M. Niu, Appl. Microbiol. Biotechnol., 2016, 100, 2011–2021.

7 S. Ge, Y. Peng, S. Wang, C. Lu, X. Cao and Y. Zhu, Bioresour. Technol., 2012, 114, 137– 143.

8 W. Li, X.-Y. Lin, J.-J. Chen, C.-Y. Cai, G. Abbas, Z.-Q. Hu, H.-P. Zhao and P. Zheng, Appl. Microbiol. Biotechnol., 2016, 100, 10203–10213.

9 Z. Si, Y. Peng, A. Yang, S. Zhang, B. Li, B. Wang and S. Wang, Environ. Sci.: Water Res. Technol., 2018, 4, 80–86.

10 L. Russ, D. R. Speth, M. S. M. Jetten, H. J. M. Op den Camp and B. Kartal, Environ. Microbiol., 2014, 16, 3487–3498.

11 K. Hanaki, Z. Hong and T. Matsuo, Water Sci. Technol., 1992, 26, 1027–1036.

12 V. Baytshtok, H. Lu, H. Park, S. Kim, R. Yu and K. Chandran, Biotechnol. Bioeng., 2009, 102, 1527–1536.

13 D. Richardson, H. Felgate, N. Watmough, A. Thomson and E. Baggs, Trends in Biotechnology, 2009, 27, 388–397.

14 J. van Rijn, Y. Tal and Y. Barak, Appl. Environ. Microbiol., 1996, 62, 2615–2620.

15 H. W. van Verseveld and A. H. Stouthamer, Arch. Microbiol., 1978, 118, 13–20.

16 J. Hinojosa, R. Riffat, S. Fink, S. Murthy, K. Selock, C. Bott, I. Takacs, P. Dold and R. Wimmer, in Proceedings of the 81st Annual Water Environment Federation Technical Exposition and Conference, Chicago, 2008, pp. 274–288.

17 T. Le, B. Peng, C. Su, A. Massoudieh, A. Torrents, A. Al-Omari, S. Murthy, B. Wett, K. Chandran, C. deBarbadillo, C. Bott and H. De Clippeleir, Water Environ. Res., 2019, 91, 1455–1465.

18 Y. Mokhayeri, R. Riffat, S. Murthy, W. Bailey, I. Takacs and C. Bott, Water Science & Technology, 2009, 60, 2485.

19 G. P. da Silva, M. Mack and J. Contiero, Biotechnol. Adv., 2009, 27, 30–39.

20 H. Lu and K. Chandran, Environ. Sci. Technol., 2010, 44, 8943–8949.

21 D. Güven, A. Dapena, B. Kartal, M. C. Schmid, B. Maas, K. van de Pas-Schoonen, S. Sozen, R. Mendez, H. J. M. Op den Camp, M. S. M. Jetten, M. Strous and I. Schmidt, Appl. Environ. Microbiol., 2005, 71, 1066–1071.

22 H. Park, A. C. Brotto, M. C. M. van Loosdrecht and K. Chandran, Water Res., 2017, 111, 265–273.

23 American Public Health Association, Standard Methods for the Examination of Water and Wastewater, American Public Health Association, American Water Works Association, Water Environment Federation, Washington, DC, 23rd edn., 2017.

24 E. M. Contreras, N. C. Bertola, L. Giannuzzi and N. E. Zaritzky, Water SA, 2002, 28, 463– 468.

25 P. D. Schloss, S. L. Westcott, T. Ryabin, J. R. Hall, M. Hartmann, E. B. Hollister, R. A. Lesniewski, B. B. Oakley, D. H. Parks, C. J. Robinson, J. W. Sahl, B. Stres, G. G. Thallinger, D. J. V. Horn and C. F. Weber, Appl. Environ. Microbiol., 2009, 75, 7537–7541.

26 S. Chen, T. Huang, Y. Zhou, Y. Han, M. Xu and J. Gu, BMC Bioinf., 2017, 18, 91–100.

27 B. J. Callahan, P. J. McMurdie, M. J. Rosen, A. W. Han, A. J. A. Johnson and S. P. Holmes, Nat. Methods, 2016, 13, 581–583.

28 J. G. Caporaso, J. Kuczynski, J. Stombaugh, K. Bittinger, F. D. Bushman, E. K. Costello, N. Fierer, A. G. Pena, J. K. Goodrich and J. I. Gordon, Nat. Methods, 2010, 7, 335–336.

29 R. Du, Y. Peng, S. Cao, S. Wang and C. Wu, Bioresour. Technol., 2015, 179, 497–504.

30 P. L. McCarty, Biotechnol. Bioeng., 2007, 97, 377–388.

31 J. C. Akunna, C. Bizeau and R. Moletta, Water Res., 1993, 27, 1303–1312.

32 H. Constantin and M. Fick, Water Res., 1997, 31, 583–589.

33 H. Sun, Q. Yang, Y. Peng, X. Shi, S. Wang and S. Zhang, Chin. J. Chem. Eng., 2009, 17, 1027–1031.

34 R. Du, S. Cao, M. Niu, B. Li, S. Wang and Y. Peng, Int. Biodeterior. Biodegrad., 2017, 122, 38–46.

35 D. Güven, Clean: Soil, Air, Water, 2009, 37, 565–573.

36 M. C. M. van Loosdrecht, M. A. Pot and J. J. Heijnen, Water Sci. Technol., 1997, 35, 41–47.

37 G. D. Drysdale, H. C. Kasan and F. Bux, Water Sci. Technol., 2001, 43, 147–154.

38 B. Liu, Y. Mao, L. Bergaust, L. R. Bakken and Å. Frostegård, Environ. Microbiol., 2013, 15, 2816–2828.

39 M. R. Betlach and J. M. Tiedje, Appl. Environ. Microbiol., 1981, 42, 1074–1084.

40 J. Dries, Water Sci. Technol., 2015, 73, 740–745.

41 L. Åmand, G. Olsson and B. Carlsson, Water Sci. Technol., 2013, 67, 2374–2398.

42 L. Gong, M. Huo, Q. Yang, J. Li, B. Ma, R. Zhu, S. Wang and Y. Peng, Bioresour. Technol., 2013, 133, 263–269.

43 Y. Z. Peng, J. F. Gao, S. Y. Wang and M. H. Sui, Water Sci. Technol., 2002, 46, 131–137.

44 C. P. L. Grady, G. T. Daigger, N. G. Love and C. D. M. Filipe, Biological Wastewater Treatment, CRC Press, Boca Raton, FL, 3rd edn., 2011.

45 H. Lu, K. Chandran and D. Stensel, Water Res., 2014, 64, 237–254.

46 M. P. Ginige, J. Keller and L. L. Blackall, Appl. Environ. Microbiol., 2005, 71, 8683–8691.

47 C. Etchebehere, I. Errazquin, E. Barrandeguy, P. Dabert and R. Moletta, FEMS Microbiol. Ecol., 2001, 35, 259–265.

48 L. Bergaust, L. R. Bakken and Å. Frostegård, Biochem. Soc. Trans., 2011, 39, 207–212.

49 R. Du, S. Cao, B. Li, M. Niu, S. Wang and Y. Peng, Water Res., 2017, 108, 46–56.

50 T. R. Thomsen, Y. Kong and P. H. Nielsen, FEMS Microbiol. Ecol., 2007, 60, 370–382.

51 C. Glass and J. Silverstein, Water Res., 1998, 32, 831–839.

