## Supporting Information for "Glycerol-driven Denitratation: Process Kinetics, Microbial Ecology, and Operational Controls"

**Calculations, Derivations, Methods**

1. Thermodynamically-derived COD requirements for denitrification using the Reaction Energetics Method for predicting bacterial yield.

2. Confirmation of assumed energy transfer efficiency factor using the Dissipation Method for predicting bacterial yield.

3. Initial reactor buffering methodology prior to pH optimization batch assays.

**List of Supplementary Tables and Figures**

Table S1. SBR feedstock components.

Figure S1. Description of feeding strategies.

Table S2. Results of Holm-Sidak post hoc multiple comparison analysis.

Table S3. Denitratation performance under different feeding strategies.

Figure S2. *Ex situ* batch assay $NO_x$ profiles at different influent COD:$NO_3^-$-N ratios.

**Calculations and Derivations**

Electron acceptor, organic electron donor, and cell synthesis half-reactions and Gibb's free energy:[1]

$$R_c:\ \ (0.20)CO_2 + (0.05)HCO_3^- + (0.05)NH_4^+ + H^+ + e^- = (0.05)C_5H_7O_2N + (0.45)H_2O$$

$$R_d:\ \ (0.21)CO_2 + H^+ + e^- = (0.07)C_3H_8O_3 + (0.21)H_2O$$

$$\Delta G_a^{0'} = 38.88\frac{kJ}{eeq}$$

$$R_{a,\ NO_3^- \to N_2}:\ \ (0.20)NO_3^- + (1.20)H^+ + e^- = (0.10)N_2 + (0.60)H_2O$$

$$\Delta G_a^{0'} = -72.20\frac{kJ}{eeq}$$

$$R_{a,\ NO_3^- \to NO_2^-}:\ \ (0.50)NO_3^- + H^+ + e^- = (0.50)NO_2^- + (0.50)H_2O$$

$$\Delta G_a^{0'} = -41.65\frac{kJ}{eeq}$$

$$R_{a,\ NO_2^- \to N_2}:\ \ (0.33)NO_2^- + (1.33)H^+ + e^- = (0.17)N_2 + (0.67)H_2O$$

$$\Delta G_a^{0'} = -92.56\frac{kJ}{eeq}$$

$$R_{in,\ acetylCoA}:\ \ (0.22)CO_2 + H^+ + e^- = (0.11)C_2H_3O + (0.33)H_2O$$ [1–3]

$$\Delta G_{in}^{0'} = 30.90\frac{kJ}{eeq}$$

$$R = f_e(R_a - R_d) + f_s(R_c - R_d) = f_eR_a + f_sR_c - R_d$$

$$1 = f_e + f_s$$

$$R_{NO_3^- \rightarrow N_2}: \ f_e[(0.20)NO_3^- + (0.07)C_3H_8O_3 + (0.20)H^+]$$
$$+ f_s[(0.05)NH_4^+ + (0.05)HCO_3^- + (0.07)C_3H_8O_3]$$
$$= f_e[(0.10)N_2 + (0.39)H_2O + (0.21)CO_2]$$
$$+ f_s[(0.05)C_5H_7O_2N + (0.24)H_2O + (0.01)CO_2]$$

$$R_{NO_3^- \rightarrow NO_2^-}: \ f_e[(0.50)NO_3^- + (0.07)C_3H_8O_3]$$
$$+ f_s[(0.05)NH_4^+ + (0.05)HCO_3^- + (0.07)C_3H_8O_3]$$
$$= f_e[(0.50)NO_2^- + (0.29)H_2O + (0.21)CO_2]$$
$$+ f_s[(0.05)C_5H_7O_2N + (0.24)H_2O + (0.01)CO_2]$$

$$R_{NO_2^- \rightarrow N_2}: \ f_e[(0.33)NO_2^- + (0.07)C_3H_8O_3 + (0.33)H^+]$$
$$+ f_s[(0.05)NH_4^+ + (0.05)HCO_3^- + (0.07)C_3H_8O_3]$$
$$= f_e[(0.17)N_2 + (0.45)H_2O + (0.21)CO_2]$$
$$+ f_s[(0.05)C_5H_7O_2N + (0.24)H_2O + (0.01)CO_2]$$

$$R_{ic,\ glycerol \rightarrow acetylCoA}: \ (0.07)C_3H_8O_3 + (0.01)CO_2 = (0.11)acetylCoA + (0.12)H_2O$$

**1. Thermodynamic derivation of COD requirements for glycerol-driven denitrification using the Reaction Energetics Method for predicting bacterial yield.**

A combination of TEEM1[1] and the modifications incorporated into the TEEM2[2] thermodynamic models was employed to determine stoichiometric coefficients for glycerol-driven chemoorganoheterotrophic denitrification.

$$\Delta G_{ic}^{0'} = \Delta G_{in}^{0'} - \Delta G_{d}^{0'} = 30.90 \frac{kJ}{eeq} - 38.88 \frac{kJ}{eeq} = -7.98 \frac{kJ}{eeq}$$

$\Delta G_{in}^{0'} = 30.90 \frac{kJ}{e^- \, eq}$ to represent the energy required to convert the cell carbon source to an intermediate compound (acetyl-CoA) prior to full oxidation.[2]

$\Delta G_d^{0'} = 38.88 \frac{kJ}{e^- eq}$ for the heterotrophic reaction.[1]

$$\Delta G_{ic}^0 = \Delta G_{ic}^{0'} - RTv_{H^+} ln[10^{-7}]$$

$$= -7.98 \frac{kJ}{eeq} - \left(0.008314 \frac{kJ}{mol \cdot K}\right)(273.15 + 23K)(0) ln[10^{-7}] =$$

$$= -7.98 \frac{kJ}{eeq}$$

$$\Delta G_{ic} = \Delta G_{ic}^0 + RTlnQ = \Delta G_{ic}^0 + RTln\left(\frac{[C_2H_3O]^{0.11}[H_2O]^{0.12}}{[CO_2]^{0.008}[C_3H_8O_3]^{0.07}}\right)$$

Assume average atmospheric $CO_{2(g)}$ concentration, $P_{CO_2} = 409ppm = 4.09 \cdot 10^{-4} atm$, therefore, $\left[CO_{2(g,headspace)}\right] = \left[CO_{2(aq)}\right] = \frac{P_{N_2}}{K_H} = \frac{4.09 \cdot 10^{-4} atm}{29.41 \frac{atm}{M}} = 1.39 \cdot 10^{-5} M$

Initial $[C_3H_8O_3] = \left(600 \frac{mgCOD}{L}\right)\left(\frac{1gCOD}{1000mgCOD}\right)\left(\frac{1molCOD}{32gCOD}\right)\left(\frac{1molC_3H_8O_3}{3.5molCOD}\right) = 5.36 \cdot 10^{-3} M$ per cycle.

Assume all glycerol is converted to acetyl-CoA, $[C_3H_8O_3] = [acetylCoA]$.

System was operated at room temperature (23°C) and buffered at $pH = 7.5$, or $[H^+] = 10^{-7.5} M$.

$$\Delta G_{ic} = -7.98 \frac{kJ}{eeq}$$

$$+ \left(0.008314 \frac{kJ}{mol \cdot K}\right)(273.15$$

$$+ 23K) ln\left(\frac{[5.36 \cdot 10^{-3} M]^{0.11}[1M]^{0.12}}{[1.39 \cdot 10^{-5} M]^{0.008}[5.36 \cdot 10^{-3} M]^{0.07}}\right) = -8.27 \frac{kJ}{eeq}$$

$$\Delta G_r^{0'} = \Delta G_a^{0'} - \Delta G_d^{0'} = -72.20 \frac{kJ}{eeq} - 38.88 \frac{kJ}{eeq} = -111.08 \frac{kJ}{eeq}$$

$$\Delta G_r^0 = \Delta G_r^{0'} - RTv_{H^+}ln[10^{-7}]$$

$$= -111.08\frac{kJ}{eeq} - \left(0.008314\frac{kJ}{mol \cdot K}\right)(273.15 + 23K)(-0.20)ln[10^{-7}]$$

$$= -119.02\frac{kJ}{eeq}$$

$$\Delta G_r = \Delta G_r^0 + RTlnQ = \Delta G_r^0 + RTln\left(\frac{[N_2]^{0.10}[CO_2]^{0.21}[H_2O]^{0.39}}{[NO_3^-]^{0.20}[C_3H_8O_3]^{0.07}[H^+]^{0.20}}\right)$$

Assume completely anoxic reactor with headspace saturated with $N_{2(g)}$, therefore,

$$[N_{2(g,headspace)}] = [N_{2(aq)}] = \frac{P_{N_2}}{K_H} = \frac{1atm}{1639.34\frac{atm}{M}} = 6.10 \cdot 10^{-4}M$$

Initial $[NO_3^-] = \left(100\frac{mgN}{L}\right)\left(\frac{1gN}{1000mgN}\right)\left(\frac{1molN}{14gN}\right) = 7.14 \cdot 10^{-3}M$ per cycle.

$$\Delta G_r = -119.02\frac{kJ}{eeq}$$

$$+ \left(0.008314\frac{kJ}{mol \cdot K}\right)(273.15$$

$$+ 23K)ln\left(\frac{[6.10 \cdot 10^{-4}M]^{0.10}[1.39 \cdot 10^{-5}M]^{0.21}[1M]^{0.39}}{[7.14 \cdot 10^{-3}M]^{0.20}[5.36 \cdot 10^{-3}M]^{0.07}[10^{-7.5}M]^{0.20}}\right) =$$

$$= -114.88\frac{kJ}{eeq}$$

$A\varepsilon\Delta G_r + \Delta G_s = 0$, at steady-state, assuming that the energy transfer efficiency from the oxidation of electron donor to capture by the electron carrier is equal to that of the electron carrier to electrons captured for cell synthesis.[1]

$$A = -\frac{\Delta G_s}{\varepsilon\Delta G_r}$$

$\Delta G_s = \frac{\Delta G_{fa} - \Delta G_d}{\varepsilon^m} + \frac{\Delta G_{in} - \Delta G_{fa}}{\varepsilon^n} + \frac{\Delta G_{pc}}{\varepsilon}$, where $\Delta G_{pc} = 18.8\frac{kJ}{eeq}$ with $NH_4^+$ as the nitrogen source for cell synthesis and $C_5H_7O_2N$ is assumed as the cell relative composition.[2] Sufficient

$NH_4^+$ was included in the feed stock for theoretical growth requirements and significant $NH_4^+$ was always remaining in the effluent indicating that additional nitrogen sources ($NO_3^-$ or $NO_2^-$) were not used for synthesis purposes as they are less energy efficient for the cell.

Since glycerol and acetyl-CoA are not C1 compounds, $\Delta G_{fa} = 0$ and $m = n$.[2]

$n = -1$ as $\Delta G_p < 0$.

$\varepsilon = 0.40$ was assumed based upon experimental data[2] and reported operational influent COD:$NO_3^-$-N ratios.[4–6]

$$\Delta G_s = \frac{\Delta G_{in} - \Delta G_d}{\varepsilon^n} + \frac{\Delta G_{pc}}{\varepsilon} = \frac{\Delta G_{ic}}{\varepsilon^n} + \frac{\Delta G_{pc}}{\varepsilon}$$

$$A = -\frac{\Delta G_s}{\varepsilon \Delta G_r} = -\frac{\frac{\Delta G_{ic}}{\varepsilon^n} + \frac{\Delta G_{pc}}{\varepsilon}}{\varepsilon \Delta G_r} = -\left( \frac{\left( \frac{-8.27 \frac{kJ}{eeq}}{0.40^{-1}} \right) + \left( \frac{18.8 \frac{kJ}{eeq}}{0.40} \right)}{(0.40)\left(-114.88 \frac{kJ}{eeq}\right)} \right) = 0.951$$

$$f_s = \frac{1}{1 + A} = \frac{1}{1 + 0.436} = 0.513$$

$$f_e = 1 - f_s = 1 - 0.513 = 0.487$$

$$R = f_e R_a + f_s R_c - R_d = (0.487) R_a + (0.513) R_c - R_d$$

$$R_{NO_3^- \to N_2}:\ NO_3^- + (0.73) C_3H_8O_3 + (0.26) NH_4^+ + (0.26) HCO_3^- + H^+$$

$$= (0.50) N_2 + (1.15) CO_2 + (0.26) C_5H_7O_2N + (3.17) H_2O$$

$$COD = (0.73\ mol\ C_3H_8O_3) \left( \frac{3.5\ mol\ O_2}{1\ mol\ C_3H_8O_3} \right) \left( \frac{32\ g\ O_2}{1\ mol\ O_2} \right) = 82.1\ g\ O_2 = 82.1\ g\ COD$$

$$NO_3^- - N = (1\ mol\ NO_3^-) \left( \frac{1\ mol\ NO_3^- - N}{1\ mol\ NO_3^-} \right) \left( \frac{14\ g\ NO_3^- - N}{1\ mol\ NO_3^- - N} \right) = 14\ g\ NO_3^- - N$$

$$COD: NO_3^- - N = 82.1\ g\ COD: 14\ g\ NO_3^- - N = 5.9: 1$$

Using this same process, assumptions of other energy-transfer efficiencies yield the following results:

| ε | 0.30 | 0.40 | 0.50 | 0.60 | 0.70 | 0.80 |
|---|---|---|---|---|---|---|
| A | 1.809 | 0.951 | 0.583 | 0.383 | 0.262 | 0.184 |
| $f_s$ | 0.356 | 0.513 | 0.632 | 0.723 | 0.792 | 0.845 |
| $f_e$ | 0.644 | 0.487 | 0.368 | 0.277 | 0.208 | 0.155 |

As can be seen, the assumption of an energy-transfer efficiency has a drastic effect and, therefore, must be confirmed.

**2. Confirmation of thermodynamic assumptions using the Dissipation Method for predicting bacterial yield.**

The Dissipation Method for predicting bacterial yield[7–9] was employed to confirm assumptions used in the thermodynamic Reaction Energetics Method determination of COD requirements to support glycerol-driven denitrification.

$\frac{D_S^{0'}}{r_{Ax}} = 200 + 18 \cdot (6 - C)^{1.8} + e^{\left[\{(3.8-\gamma_D)^2\}^{0.16} \cdot (3.6+0.4C)\right]}$, which describes the heat (Gibbs free energy) dissipated during growth or production of 1 C-mole of biomass.

$C = 3$, which represents the number of carbon atoms in a mole of glycerol.

$\gamma_D = 4.667$, degree of reductance of the carbon in glycerol as the electron donor.[7]

$$\frac{D_S^{0'}}{r_{Ax}} = 200 + 18 \cdot (6 - 3)^{1.8} + e^{\left[\{(3.8-4.667)^2\}^{0.16} \cdot \{3.6+(0.4)(3)\}\right]} = 428.06 \frac{kJ}{c\,mol}$$

$Y_{DX} = \frac{\gamma_D}{\gamma_X} \frac{\Delta G_{eD}^{0'} - \Delta G_{eA}^{0'}}{\left(\Delta G_{eD}^{0'} - \Delta G_{eA}^{0'}\right) + \left[\left(\frac{D_S^{0'}}{r_{Ax}} \cdot \frac{1}{\gamma_X}\right) + \left(\Delta G_{eX}^{0'} - \Delta G_{eD}^{0'}\right)\right]}$, which represents the bacterial cell yield on the electron donor.

$\Delta G_{eD}^{0'} = 38.88 \frac{kJ}{eeq}$, Gibbs standard free energy for glycerol as the electron donor.[1]

$\Delta G_{eA}^{0'} = -72.20 \frac{kJ}{eeq}$, Gibbs standard free energy for $NO_3^-$ as the electron acceptor.[1]

$\Delta G_{eX}^{0'} = 38.80 \frac{kJ}{eeq}$, assuming $\Delta G_{fX}^{0'} = -67 \frac{kJ}{c{-}mol}$.[10]

$\gamma_X = 4.2$, degree of reductance of the carbon in biomass.[7]

$$Y_{DX} = \left(\frac{4.667}{4.2}\right)\left[\frac{38.88\frac{kJ}{eeq} - \left(-72.20\frac{kJ}{eeq}\right)}{\left\{38.88\frac{kJ}{eeq} - \left(-72.20\frac{kJ}{eeq}\right)\right\} + \left\{\left(428.06\frac{kJ}{c\ mol} \cdot \frac{1}{4.2}\right) + \left(38.80\frac{kJ}{eeq} - 38.88\frac{kJ}{eeq}\right)\right\}}\right] = 0.580 \frac{c\ mol_X}{c\ mol_D}$$

$$Y_{DX} = 0.522 \frac{eeq_X}{eeq_D}$$

In terms of eeq, $Y_{DX} = f_s^0$, therefore, $f_s^0 = 0.522 \frac{eeq_X}{eeq_D}$.

$f_e^0 = 1 - f_s^0 = 1 - 0.522 = 0.487$

Comparison of $f_s^0$ calculated using the Dissipation Method with $f_s$ calculated using the Reaction Energetics Method indicates that the energy-transfer efficiency, ε, inherent in the Dissipation Method calculations is $\varepsilon = 0.406$. This confirms the validity of the assumption of $\varepsilon = 0.40$ in the Reaction Energetics Method calculations.

While these calculations are at standard state, it has been shown that there is little difference between predictions at standard state and non-standard state in certain instances provided system pH is close to neutral, substrate concentrations are low, and $\Delta G_{eD}^{0'} - \Delta G_{eA}^{0'} > 20 \frac{kJ}{eeq}$.[2,9]

Additionally, as they are simply being used to confirm assumptions made using the reaction energetics method, calculations were not made to convert to non-standard state conditions.

**3. Initial reactor buffering methodology prior to pH optimization batch assays.**

For pH optimization batch assays, the medium was initially buffered to approximately pH 9.0 but left unbuffered for the remainder of each experiment during which the pH ranged from 7.2 to 9.0.

***Table S1.*** *Components of SBR feed including trace nutrients. *Trace nutrients were dissolved in deionized water.*

| **SBR Feed** (mg per 100 L SBR feed) | |
|---|---|
| $NO_3^-$-N | 10,000.0 |
| $NH_4^+$-N | 2,500.0 |
| $MgSO_4 \cdot 7H_2O$ | 20,000.0 |
| $KH_2PO_4$ | 8,700.0 |
| $CaCl_2 \cdot 2H_2O$ | 2,000.0 |
| NaOH | for pH adjustment |
| **Trace Nutrients*** (mg per 100 L SBR feed) | |
| EDTA·$Na_2$ | 2,010.1 |
| $FeSO_4 \cdot 7H_2O$ | 500.4 |
| $MnCl_2 \cdot 4H_2O$ | 172.2 |
| $ZnSO_4 \cdot 7H_2O$ | 43.1 |
| $CuSO_4 \cdot 5H_2O$ | 25.0 |
| $CoCl_2 \cdot 6H_2O$ | 23.8 |
| $Na_2MoO_4 \cdot 2H_2O$ | 10.0 |
| $NiSO_4 \cdot 6H_2O$ | 2.1 |
| $H_3BO_3$ | 1.1 |

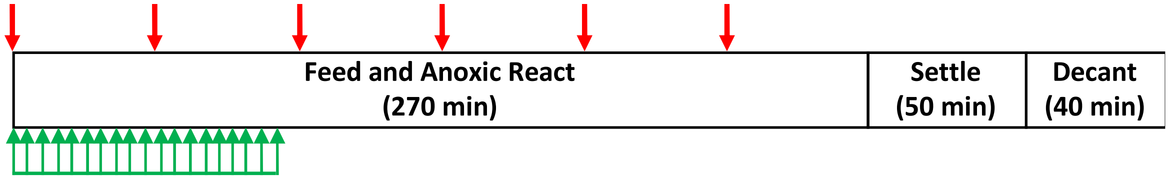


***Figure S1.*** *Two feeding strategies, semi-continuous (green arrows; 75-min $NO_3^-$ feed with concurrent 72-min glycerol feed, influent COD:$NO_3^-$-N=2.4:1) and pulse (red arrows; each pulse contained 4-min $NO_3^-$ feed with concurrent 1-min glycerol feed, influent COD:$NO_3^-$-N=2.4:1), were investigated to determine their impact on $NO_2^-$ accumulation.*

***Table S2.*** *Results of Holm-Sidak post hoc multiple comparison analysis to determine between which NARs a significant difference exists (statistical significance exists at $p<0.05$ and is demarcated using bold font).*

| Inf. COD:$NO_3^-$-N | 2.5 ($\bar{x}$=0.65) | 2.8 ($\bar{x}$=0.69) | 3.0 ($\bar{x}$=0.62) | 4.0 ($\bar{x}$=0.57) | 5.0 ($\bar{x}$=0.11) |
|---|---|---|---|---|---|
| 2.5 | | 0.496 | 0.755 | 0.319 | **0.000** |
| 2.8 | | | 0.147 | **0.006** | **0.000** |
| 3.0 | | | | 0.329 | **0.000** |
| 4.0 | | | | | **0.000** |

***Table S3.*** *Denitratation performance under continuous and pulse operational feeding strategies.*

| Influent $COD:NO_3^-$-N | SRT [d] | Operational Feeding Strategy | Avg $NO_{3,eff}^-$ [mg/L $NO_3^-$-N] | Avg $NO_{2,eff}^-$ [mg/L $NO_2^-$-N] |
|---|---|---|---|---|
| 2.4 | 3 | Pulse $NO_3^-$ Pulse COD | 11.3 ± 3.3 | 86.4 ± 7.5 |
| | | Continuous $NO_3^-$ Continuous COD | 16.0 ± 5.5 | 70.1 ± 8.4 |

Contrary to the continuous operational feeding strategy, the pulse operational feeding strategy reduced nearly 90% of the influent $NO_3^-$ despite the limited reaction time for late occurring pulses of $NO_3^-$ and glycerol indicating that influent $NO_3^-$ underwent rapid reduction upon entering the system. This observation was consistent with other studies which reported that specific denitrification rates are higher for pulse-type feeding strategies as compared to continuous feeding strategies resulting in a faster reduction of influent $NO_3^-$.[11,12] Martins et al.[11] determined that maximum specific denitrification rates were considerably lower for SBR systems with long feeding periods that mimicked continuously-fed, completely mixed systems, than in plug flow-type systems. Similarly, Ryu et al.[12] found that denitrification rates were fastest during slug feeding followed in order by intermittent and continuous feeding strategies during their evaluation of fermented food waste as an external carbon source for nutrient removal in an SBR.

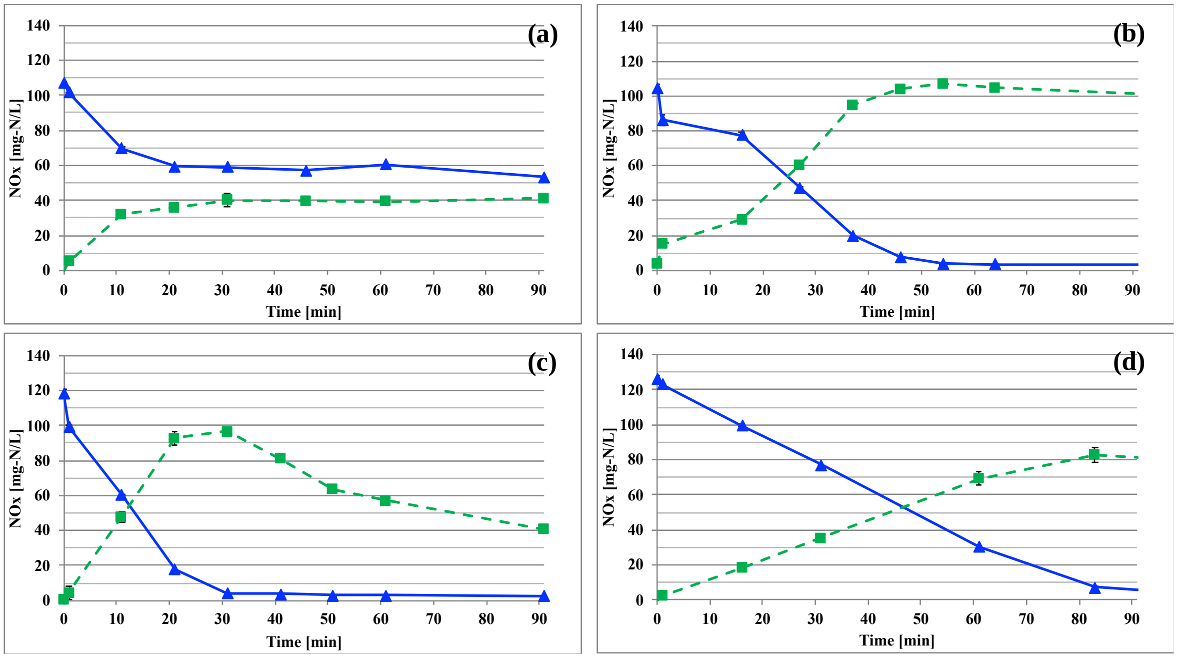


***Figure S2.*** *Representative* ex situ *$NO_3^-$-N (▲, solid line) and $NO_2^-$-N (■, dotted line) profiles at influent COD:$NO_3^-$-N ratios (a) 2.5, (b) 3.0, (c) 5.0, (d) 10.0.* Ex situ *batch assays were performed using biomass acclimated at each influent COD:$NO_3^-$-N ratio in the parent reactor for at least four SRTs.*

### References


1 B. E. Rittmann and P. L. McCarty, *Environmental Biotechnology: Principles and Applications*, McGraw-Hill, Boston, 2001.

2 P. L. McCarty, *Biotechnol. Bioeng.*, 2007, **97**, 377–388.

3 R. K. Thauer, K. Jungermann and K. Decker, *Bacteriol. Rev.*, 1977, **41**, 100–180.

4 J. Hinojosa, R. Riffat, S. Fink, S. Murthy, K. Selock, C. Bott, I. Takacs, P. Dold and R. Wimmer, in *Proceedings of the 81st Annual Water Environment Federation Technical Exposition and Conference*, Chicago, 2008, pp. 274–288.

5 H. Lu and K. Chandran, *Environ. Sci. Technol.*, 2010, **44**, 8943–8949.

6 J. C. Akunna, C. Bizeau and R. Moletta, *Water Res.*, 1993, **27**, 1303–1312.

7 J. M. VanBriesen, *Biodegradation*, 2002, **13**, 171–190.

8 J. J. Heijnen and J. P. van Dijken, *Biotechnol. Bioeng.*, 1992, **39**, 833–858.

9 J. J. Heijnen, M. C. M. van Loosdrecht and L. Tijhuis, *Biotechnol. Bioeng.*, 1992, **40**, 1139–1154.

10 J. A. Roels, *Biotechnol. Bioeng.*, 1980, **22**, 2457–2514.

11 A. M. P. Martins, J. J. Heijnen and M. C. M. van Loosdrecht, *Biotechnol. Bioeng.*, 2004, **86**, 125–135.

12 H.-D. Ryu, S.-I. Lee, K.-Y. Chung and K.-Y. Kim, *Environ. Eng. Sci.*, 2007, **24**, 1467–1474.
